# The isotopic nitrogen turnover rate as a proxy to evaluate in the long-term the protein turnover in growing ruminants

**DOI:** 10.1101/543116

**Authors:** Gonzalo Cantalapiedra-Hijar, Hélène Fouillet, Céline Chantelauze, Nadezda Khodorova, Lahlou Bahloul, Isabelle Ortigues-Marty

**Affiliations:** Université Clermont Auvergne, INRA, VetAgro Sup, UMR Herbivores, F-63122 Saint-Genès-Champanelle, France; UMR Nutrition Physiology and Ingestive Behavior, AgroParisTech, INRA, Paris-Saclay University, F-75005 Paris, France; Adisseo France S.A.S., Antony, France

**Keywords:** protein turnover, isotope, ^15^N, ruminant, beef cattle

## Abstract

Protein turnover is an energy-consuming process essential for ensuring the maintenance of living organisms. Gold standard methods for protein turnover measurement have inherent drawbacks precluding their generalization for large farm animals and during long periods. We proposed here a non-invasive proxy of the whole-body fractional protein degradation rate (**WBPT**; whole-body protein turnover for a growing animal) in the long term and in a large number of beef cattle. The proxy is based on the rate at which urine-N and plasma proteins are progressively depleted in ^15^N after a slight decrease in the isotopic N composition of diet (i.e. diet-switch). We aimed to test the ability of this proxy to adequately discriminate the WBPT of 36 growing-fattening young bulls assigned to different dietary treatments known to impact the protein turnover rate: the protein content (Normal vs High) and amino acid profile (balanced vs unbalanced in methionine). The ^15^N depletion rate found in plasma proteins represented their fractional synthesis rate, whereas the slowest depletion rate found in urines was interpreted as a proxy of the WBPT. The proxy here tested in urines suggested different WBPT values between Normal vs High protein diets but not between balanced vs unbalanced methionine diets. In contrast, the proxy tested in plasma indicated that both dietary conditions affected the fractional synthesis rate of plasma proteins. We consider that the rate at which urines are progressively ^15^N-depleted following an isotopic diet-switch could be proposed as a non-invasive proxy of the whole-body protein turnover rate in large farm animals.

## Introduction

Protein turnover refers to the renewal of proteins in a cell, tissue or organism and involves the continuous degradation and synthesis of proteins not leading to net changes in protein mass. The protein accretion occurring in the growing animal is the result of a greater intensity of protein synthesis over protein degradation. In the growing state, the protein turnover rate (amount of replaced proteins per unit of time) equals to total protein degradation or alternatively to the amount of freshly synthesized protein necessary to replace the degraded protein. The protein turnover in growing ruminants is huge compared to the net protein deposition with as high as 94% of the whole body protein synthesis only serving to counterbalance the protein degradation occurring in fattening young bulls (Lobley, 2003). Besides, this huge turnover is accompanied by a high-energy cost for the ruminant, with only protein synthesis accounting for 23% of total energy expenditure in ruminants (Caton et al., 2000). Nevertheless, this dynamic state represents an essential mechanism for life by enabling maintenance services such as metabolic regulation, cellular repair and rapid adaptation against environmental changes, among other functions. This double faceted role of protein turnover (“service” function for maintenance vs energy-consuming mechanism) explain why this animal trait has been of interest for animal nutritionist and physiologist in the last decades.

Standard methods for whole-body protein turnover assessment are based on either precursor or end-product methods (Waterlow, 1984) consisting in continuous or single dose intravenous infusions of stable isotope labelled amino acids (**AA**) and the quantification of the labelling in either the free AA precursor pool (plasma) or catabolic end-products (urine), respectively. The main drawbacks associated to the evaluation of protein turnover rate through these standard methods are i) short-term evaluation (from few hours [precursor method] to few days [end-product method]), ii) invasive procedure and usually non-physiological conditions, iii) the requirement of a controlled environment with confined animals and usually steady-state conditions [precursor method] iv) expensive methods which precludes applying it to a big number of large farm animals such as beef cattle. Alternative, less expensive and less invasive methods have been explored, such as total urinary 3-methyl-histidine excretion, to evaluate the whole-body protein degradation. However, 3-methyl-histidine urinary excretion is also based on relatively short term evaluation (i.e. usually between 4 and 6 days), restricted to myofibrillar muscle protein (myosin-actin) turnover, not applicable for sheep or pigs (Nishizawa et al., 1979) and subjected to an accurate estimation of the muscle mass in the body (Castro-Bulle et al., 2007).

A new isotopic approach was originally developed in animal ecophysiology (Fry and Arnold, 1982; Tieszen et al., 1983) to model the rate at which animal’s tissues incorporate the isotopic signature of a new diet in view of accurate food web assessment. It is currently being applied mainly in marine animal species (Abimorad et al., 2014; Mohan et al., 2016) to evaluate the metabolic turnover rate of animal’s tissues. This isotopic method consists in following the dynamic of isotopic change (i.e. enrichment or depletion) of animal’s tissues immediately after a change in the natural isotopic composition of the diet (i.e. diet-switch). The rate at which animal’s tissues will incorporate the isotopic signature of the new diet seems to be mostly driven by their protein turnover rate (Carleton and Martinez del Rio, 2005; MacAvoy et al., 2005; Braun et al., 2013) and some authors suggest that protein turnover rate might be predicted from isotopic incorporation studies (Carleton et al., 2008). This approach has already been explored in ruminant’s tissues (Harrison et al., 2011; Bahar et al., 2014) but, as its original purpose, just for assessing the time span needed for muscles to reflect the isotopic signature of the diet in view of meat authentication (i.e isotopic dietary reconstruction). However, the isotopic approach used in those studies in ruminant and marine animal species did not involve repeated measures from the same animal, but only one tissue sample per animal, and individuals slaughtered at different times. Thus, these studies only informed about the turnover of specific sampled tissues and not of the whole animal body. Because most of the nitrogen from animal proteins is excreted through urines, once degraded and their AA catabolized, our hypothesis is that the urinary isotopic N turnover rate assessed from multi-point sampling from the same animal is a non-invasive proxy of the long term whole-body fractional protein degradation rate (**WBPT**; whole-body protein turnover in a growing animal).

Thus, the aim of this study was to adapt the tissue isotopic turnover approach (MacAvoy et al., 2005; Abimorad et al., 2014) to the ruminant’s urine and test it as a non-invasive, less disturbing proxy of the long term WBPT in a relatively large number of growing-fattening beef cattle. In addition, we also measured the isotopic turnover of plasma proteins as an indicator of the fractional protein synthesis rate (**FSR**) of plasma proteins. Because no gold standard method is currently available for measuring the WBPT in the long term (several months) we evaluated this proxy by assessing its ability to detect differences across two dietary factors known to impact the whole-body protein turnover in a different extent: i) the protein content and intake as a strong and well-known effect (Waterlow, 2006) and ii) the dietary amino acid profile as a less important modulator of protein turnover rate in cattle (Wessels et al., 1997). For this purpose, we measured the kinetics of nitrogen isotopic (δ^15^N) depletion in urines and plasma proteins over a 5-month period following a slight decrease in the δ^15^N of the diet in 36 fattening young bulls fed diets formulated at two different protein and methionine contents. Preliminary results has been published as an abstract (Cantalapiedra-Hijar et al., 2017).

## Material and methods

The experiment was conducted at Herbipôle (Inra, UE 1414, Theix, France) in compliance with the National Legislation on Animal Care. The C2EA-02 animal research ethics committee (Auvergne, France) prospectively approved this research and thereafter the Ministry of Agriculture (France) validated it with the approval number #7180-2016101016361277v4.

### Animals, diets and performance test

Thirty-six growing-fattening Charolais bulls (320±33kg and 266±22d) were assigned to one of the four experimental diets (n=9/treatment) resulting from a factorial 2 × 2 design: two dietary metabolizable protein levels (100% [Normal] vs 120% [High] of requirements; INRA, 2018) crossed with two dietary methionine content (unbalanced diet [2.0 g Met/100 g metabolizable protein] vs balanced diet [2.6 g Met/100 g metabolizable protein]). All diets were iso-NE per kg DM and consisted of around 54% grass silage, 6% of wheat straw and 40% concentrate with amounts adjusted daily to ensure at least 10% of refusals.

### Dietary isotopic N switch

The method here applied to assess the WBPT was based on the rate at which tissues and animal N pools incorporate a new dietary ^15^N isotopic signature following a dietary isotopic switch (Carleton and Martinez del Rio, 2005). However, because the range of natural ^15^N abundance values (**δ**^**15**^**N**; ^15^N/^14^N ratio deviation from the international standard [atmospheric N_2_]) in ruminant’s feed remains quite narrow (between 0 and 5‰ on average according to our experience [Cantalapiedra-Hijar et al., 2015]) we decided to create an artificial dietary isotopic switch by incorporating in the diet extrinsic ^15^N labelled urea as previously reported (Bahar et al., 2014). The ^15^N-urea once degraded into ^15^N-ammonia and taken up by the rumen microorganisms will enrich the animal’s tissues through the absorbed microbial ^15^N-AA. Results from a preliminary trial aiming to define the conditions for implementing this method in ruminants showed that for a given animal the fraction of ^15^N-urea bypassing directly to the urines, without being first incorporated into microbial proteins, was highly variable across time and individuals and led to problems for fitting the progressive urinary ^15^N-enrichment post-diet switch (data not shown). Thus, we decided to evaluate the animal ^15^N turnover rates during a depletion (Abimorad et al., 2014) rather than enrichment (Bahar et al., 2014) phase. For this, animals were progressively enriched in ^15^N over 35 days while adapting to their respective diet; each animal received daily in the morning (0830) a capsule (10 × 3mm) containing 20 mg of ^15^N labelled urea (98% APE; Sigma–Aldrich, St. Louis, USA). This amount of ^15^N labelled urea and time was chosen for enhancing the urinary ^15^N between 10 and 15‰ above that of the basal diet according to results obtained from a preliminary trial. The capsule was mixed with around 200 grams of concentrate (the one assigned to the animal) in a small bucket within the feeder just before the only meal distribution (0900). Twenty minutes after administration it was systematically checked that the capsule had been swallowed by the animal. Less than 5% of capsules were administered by hand directly in the mouth when animals did not swallow it within the first twenty minutes. The rest of the diet was then distributed to the feeder. On day 36, animals no longer received the ^15^N labelled urea, and blood and urine were sampled from that day (d0) onwards.

### Sampling and ^15^N analysis

Blood was sampled at 0900 by venipuncture from the caudal vein of all animals on d0, d3, d7, d11, d15, d21, d35, d49, d78 and d141 (n=10 per animal) after stopping the ^15^N labelled urea administration (d0 refers to 24h after the last ^15^N-urea administration). Blood was collected into 9 mL evacuated tubes (BD vacutainer, Plymouth, UK) containing lithium heparin as an anticoagulant, centrifuged within the first hour at 2500 g for 15 min at 4°C and stored at - 20°C for determination of δ^15^N values in total plasma proteins as previously described (Cantalapiedra-Hijar et al., 2015). Urinary spot samples were obtained from all animals between 0900 and 1000 on d0, d1, d2, d3, d4, d7, d9, d11, d14, d17, d35, d70, d142 (n = 13). For urinary sampling, animals were head blocked during 1h while eating in the morning and a bucket (30 × 30 × 15 cm) placed around the penis through two elastic ropes knotted on the back of the animal. Most of animals urinated during the first hour and only few samplings were conducted beyond the first hour and always within the first three hours. Twenty mL of urine were then transferred into one tube containing 1 mL of 30%H_2_SO_4_ and filtered later in the lab through a 30 µm standard filter paper to remove fine particles. Filtered and acidified spot urine samples were stored at −20°C before determination of δ^15^N values. Freeze-dried plasma proteins and liquid urinary samples pipetted onto nitrogen-free absorbent (chromosorb) were weighted in tin capsules and analyzed for N stable isotope composition (δ^15^N) by using an isotope-ratio mass spectrometer (Isoprime, VG Instruments, Manchester, UK) coupled to an elemental analyzer (EA Vario Micro Cube, Elementar, Germany). Tyrosine was used as internal standard and included in every run to correct for possible variations in the raw values determined by the mass spectrometer. Results were expressed using the delta notation.

### Isotopic turnover rate modelling and statistical analysis

The post-diet switch δ^15^N kinetics measured in each pool (plasma protein and urine) were carefully analyzed according to Martinez del Rio and Carleton (2012) by testing if they obey to 1st-order or higher-order (2^nd^-order) kinetics according to the following mono- and bi-exponential models, respectively:

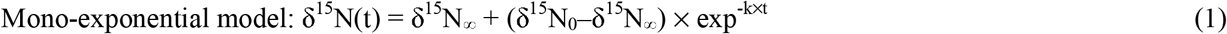

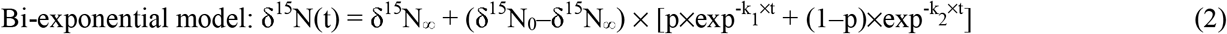

where t (d) is the time since the ^15^N diet-switch, δ^15^N(t) (‰) is the pool δ^15^N value at time t, δ^15^N_0_ (‰) is the pool initial δ^15^N value, and δ^15^N_∞_(‰) is the asymptotic value of the pool after the animal has reached isotopic steady state with its basal diet (without ^15^N-urea administration). In the mono-exponential model, k (d^−1^) is the fractional isotopic turnover rate of the pool, while in the bi-exponential model, k_1_ and k_2_ (d^−1^) are two distinct fractional isotopic turnover rates and p and (1-p) t their respective contributions to the whole isotopic turnover.

To diagnose if a mono-exponential model was sufficient to adequately fit the δ^15^N kinetics, we used the reaction progress variable approach ((Martinez del Rio and Carleton, 2012; Cerling et al., 2007) that it is based on the rearrangement of equation [1] to yield:

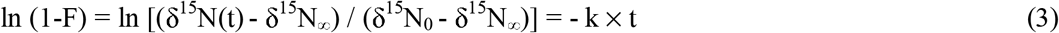

where (1-F) measures the remaining δ^15^N distance to the new equilibrium as a proportion of the total isotopic distance between the initial and the asymptotic δ^15^N reached at equilibrium. As illustrated in S1 Appendix, we decided what model was required on the basis of a visual inspection of a plot of ln(1-F) against time, depending on whether ln(1-F) was a decreasing linear function of time with slope equal to -k (mono-exponential model) or a sequence of 2 lines of increasingly shallow slopes equal to –k_1_ and –k_2_ (bi-exponential model). This graphical analysis was further supported by the Akaike Information criterion, where lower values indicates a superiority of one model over another.

The model parameters (i.e., δ^15^N_∞_, δ^15^N_0_ and k for the mono-exponential model, and δ^15^N_∞_, δ^15^N_0_, k_1_, k_2_ and p for the bi-exponential model) were estimated using a non-linear procedure and statistically analyzed through a non-linear mixed-effect model (nlme package; Pinheiro and Bates, 2000) fitted by maximum likelihood in the R software with experimental dietary factors (protein content, methionine content and their interaction) considered as fixed effects and the animal as a random effect. To avoid over-parametrization in the most complex model (bi-exponential) and avoid a failure for the model to converge, the random effect of animal was only considered on those parameters where significant animal variability was noted through graphical and statistical analysis (Pinheiro and Bates, 2000). Significant effects were declared when the *P*≤0.05 and a trend was considered when 0.05>*P*<0.10.

## Results

No significant interaction (*P* > 0.05) between protein level and methionine concentration was observed for any of the variables here analyzed. Thus, for the shake of clarity, results are presented independently for each of these two experimental factors. During the length of the applied approach (around 5 months) bulls fed High protein diets had on average higher (*P* = 0.005) daily gains (1.82 kg/d) compared to those fed Normal protein diets (1.63 kg/d on average). On the other hand, methionine balanced diets tended (*P* = 0.09) on average to promote higher daily gains (1.78 kg/d) than bulls fed unbalanced diets (1.67 kg/d).

### Urinary isotopic turnover rate

For δ^15^N kinetics in urines, the reaction progress approach clearly identified two independent slopes with a cut-off between d4 and d7 (Insets in Figs 1 and 2). This demonstrated the existence of two distinct rates, a rapid and a slow, of urinary δ^15^N depletion after the diet-switch, and justified the need of a bi-exponential model to adequately fit these data. The individual δ^15^N kinetics were indeed correctly fitted for almost all animals (r2 ≥ 0.96; n=34) using a bi-exponential model, except for 2 animals that were badly fitted with no apparent explanation and that were therefore excluded from the analyses. When all data were pooled and analyzed through a mixed-effect bi-exponential model, only a significant effect of the dietary protein content was observed on the fractional rates of ^15^N-depletion in urines, where animals fed High protein diets showed higher values during both the first rapid (89.9 vs. 70.0 %/d; *P* = 0.008) and the second slow (10.3 vs. 8.01%/d; *P* < 0.001) phases of δ^15^N turnover compared to animals fed Normal protein diets (Fig 1). No effect of methionine content was observed on any model parameter (*P* > 0.10; Fig 2).

**Figure 1.**
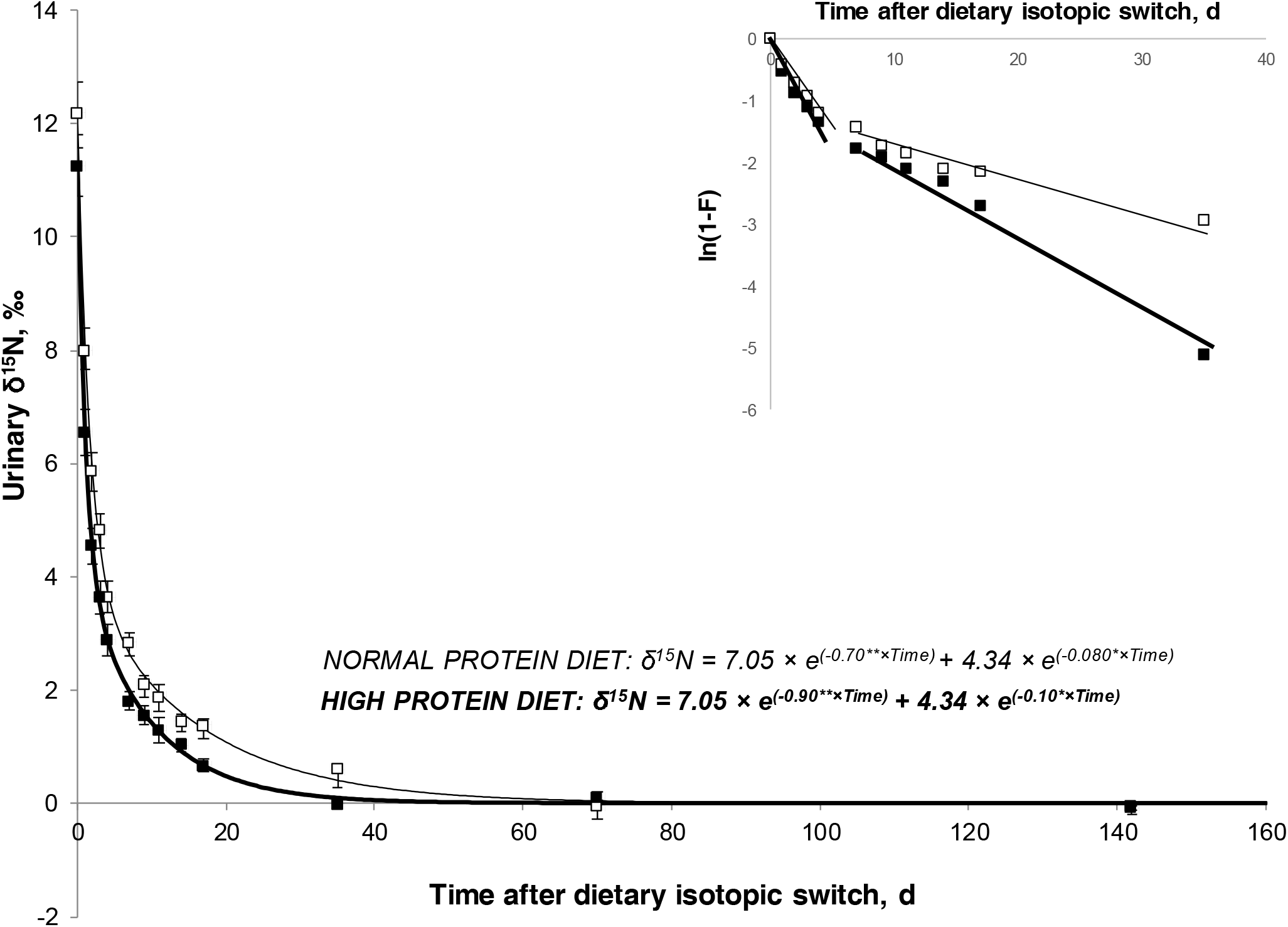
Kinetics of ^15^N-depletion in urines after a ^15^N diet-switch (see Material and Methods) in Charolais fattening bulls fed either Normal (thin line) or High (thick line) protein diets. Compared to animals fed Normal protein diet (n = 17), animals fed High protein diets (n = 17) showed higher fractional rates of urinary ^15^N-depletion during both the first rapid (89.9 vs. 70.0%/d; P = 0.008) and the second slow (10.3 vs. 8.01 %/d; P < 0.001) phases. Inset represents the reaction-progress variable approach [ln(1-F); see Material and Methods] diagnosing that adequately fitting the ^15^N-depletion kinetics in urine required a bi-exponential model with two rates for two - rapid and slow - phases (Martinez del Rio and Carleton, 2012). Significant differences in model parameters across treatments are depicted by stars (\*\**P* < 0.01; \**P* < 0.05).

**Figure 2.**
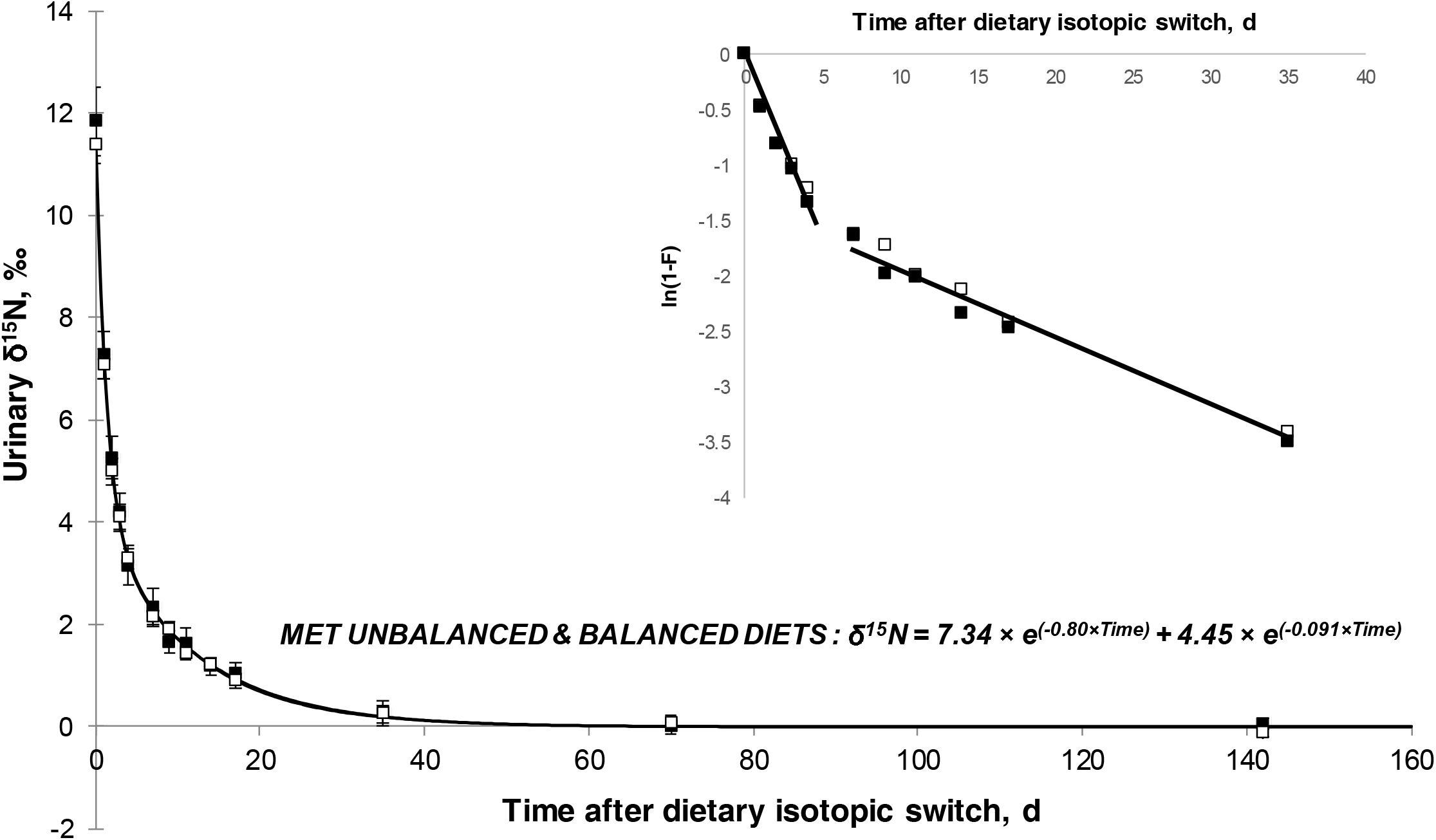
Kinetics of ^15^N-depletion in urines after a ^15^N diet-switch (see Material and Methods) in Charolais fattening bulls fed either balanced (thin line) or unbalanced (thick line) diets in terms of methionine content. Compared to animals fed diets unbalanced in methionine (n = 17), animals fed diets balanced in methionine (n = 17) showed similar fractional rates of urinary ^15^N-depletion during both the first rapid (80.0%/d; *P* = 0.92) and the second slow (9.10%/d; *P* = 0.80) phases. Inset represents the reaction-progress variable approach [ln(1-F); see Material and Methods] diagnosing that adequately fitting the ^15^N depletion kinetics in urine required a bi-exponential model with two rates for two - rapid and slow – phases (Martinez del Rio and Carleton, 2012).

### Plasma isotopic turnover rate

For δ^15^N kinetics in plasma proteins, the reaction progress approach showed a single slope (Insets in Figs 3 and 4). This demonstrated the existence of a single, homogenous rate of δ^15^N depletion in plasma proteins after the diet-switch, and justified that a mono-exponential model was sufficient to adequately fit these data. The individual δ^15^N kinetics in plasma proteins were indeed correctly fitted (r2 ≥ 0.98; n=36) through a mono-exponential asymptotic model. When all data were pooled and analyzed through a mixed-effect asymptotic model, lower plasma protein δ^15^N values were observed in animals fed High compared to Normal protein diets (Fig 3) on first day (d0; *P* = 0.04) and at equilibrium (d142; *P* = 0.09). In contrast, the effect of balancing diets in terms of methionine content tended (*P* = 0.09) to have a higher δ^15^N value in plasma proteins on d0 (Figure 4) but not at equilibrium (*P* = 0.47). The fractional ^15^N-depletion rate in plasma proteins was higher in animals fed High compared to Normal protein diets (4.42 vs 4.08%/d; *P* = 0.02) and in those fed diets balanced compared to unbalanced in methionine (4.38 vs 4.10%/d; *P* = 0.05).

**Figure 3.**
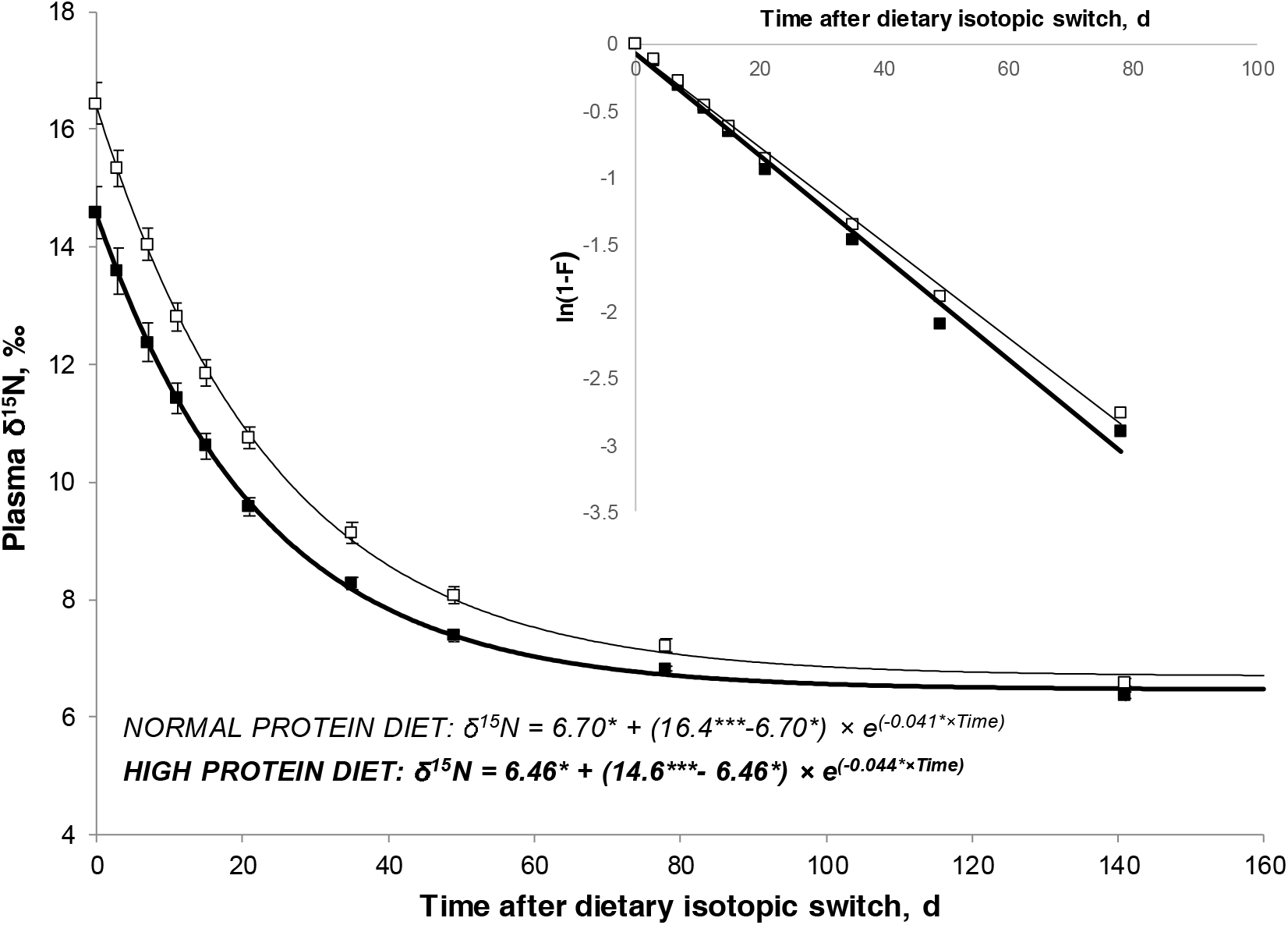
Kinetics of ^15^N depletion in plasma proteins after a ^15^N diet switch (see Material and Methods) in Charolais fattening bulls fed either Normal or High protein diets. Compared to animals fed Normal protein diets (n = 18), animals fed High protein diets (n = 18) showed lower plasma protein δ^15^N values at time 0 (14.6 vs. 16.4‰; P < 0.001) and higher fractional rate of plasma protein ^15^N-depletion (4.42 vs. 4.08 %/d; *P* = 0.02). Inset represents the reaction-progress variable approach [ln(1-F); see Material and Methods] diagnosing that a mono-exponential model was sufficient to adequatly fit the ^15^N-depletion kinetics in plasma proteins (Martinez del Rio and Carleton, 2012). Significant differences in model parameters across treatments are depicted by stars (\*\*\**P* < 0.001; \*\**P* < 0.01; \**P* < 0.05).

**Figure 4.**
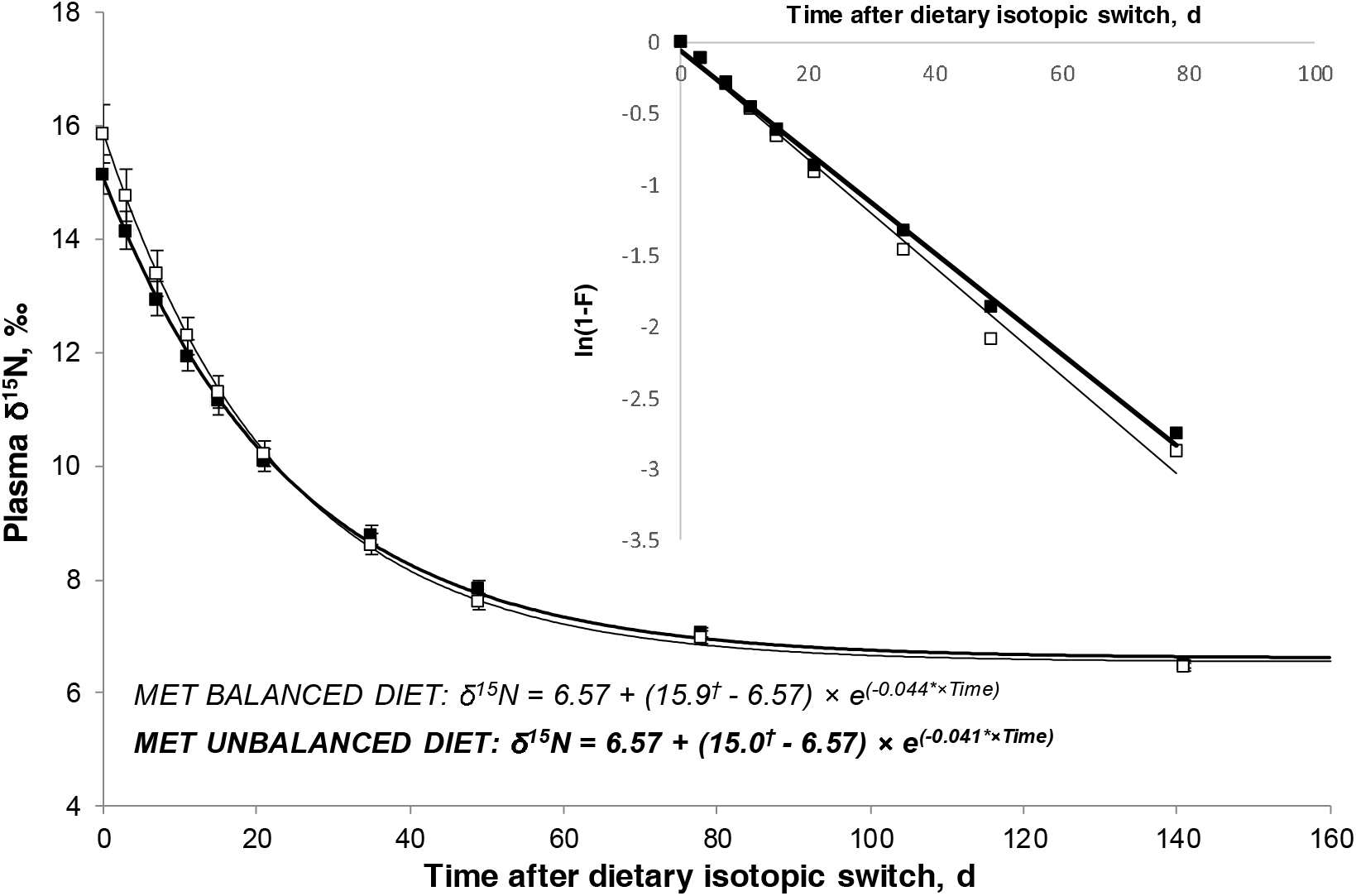
Kinetics of ^15^N depletion in plasma proteins after a ^15^N diet switch (see Material and Methods) in Charolais fattening bulls fed either balanced or unbalanced diets in terms of methionine content. Compared to animals fed diets unbalanced in methionine (n = 18), animals fed diets balanced in methionine (n = 18) showed higher plasma protein δ^15^N values at time 0 (15.9 vs. 15.0‰; *P* = 0.05) and higher fractional rate of plasma protein ^15^N-depletion (4.38 vs. 4.10%/d; *P* = 0.05). Inset represents the reaction-progress variable approach [ln(1-F); see Material and Methods] diagnosing that a mono-exponential model was sufficient to adequatly fit the ^15^N depletion kinetics in plasma proteins (Martinez del Rio and Carleton, 2012). Significant differences in model parameters across treatments are depicted by symbols (^†^*P* < 0.10; \**P* < 0.05).

## Discussion

Following a diet change, animal’s tissues progressively assimilate the isotopic signature of the new diet (Carleton and Martinez del Rio, 2005). The rate at which this occur is known as the isotopic turnover rate and depends largely on how fast the metabolic tissue replacement occurs (Tieszen et al., 1983; Arneson et al., 2006). We adapted the tissue isotopic turnover approach, extensively explored by ecologists (Martinez del Rio and Carleton, 2012), to the case of ruminant’s urines and plasma proteins to respectively evaluate in the long term (5 months) the WBPT (protein turnover in a growing animal) and the FSR of plasma protein in 36 growing-fattening cattle. This isotopic approach is simple to set up, relatively cheap (estimated to be less than 300 dollars per bull), allowing to obtain measurements over long periods (several months) and without perturbing animals (unlike classic tracer methods with invasive procedures under non-physiological conditions). Although the mechanistic interpretation of isotopic turnover rates is always a complex issue (Martinez del Rio and Anderson-Sprecher, 2008), it is proposed that the rates at which urines and plasma proteins are progressively ^15^N-depleted following an isotopic diet-switch represent non-invasive (or less invasive) proxies of the long-term WBPT and plasma protein FSR, respectively. Such proxies could be very useful for future studies on feed efficiency (protein turnover as an energy-consuming process), animal robustness (protein turnover as a maintenance service) and meat quality (in vivo protein turnover associated to the rate of post mortem proteolysis and thus to meat tenderness) carried out on large number of animals.

### Biological meaning of isotopic turnover rates

The simplicity of the method here described may contrast with the need to address some methodological considerations for interpreting our results. The rate at which animal’s tissues will incorporate the isotopic signature of the new diet seems to be mostly driven by their protein turnover rate according to many previous reports (Carleton and Martinez del Rio, 2005; MacAvoy et al., 2005; Braun et al., 2013). Moreover, this has been confirmed by mechanistic models (Martinez del Rio and Carleton, 2012; Poupin et al., 2012), demonstrating that the main driver determining the rate of assimilation of the new dietary δ^15^N value into a tissue after a diet-switch is the protein FSR of that tissue. Thus, the isotopic turnover rate we found in the plasma protein pool (i.e., the k model parameter; Figs 3 and 4) represented its own FSR (i.e., the sum of its fractional degradation rate and fractional growth rate) and may be proposed as a way to indirectly evaluate the liver FSR of plasma proteins. In contrast, to the best of our knowledge the isotopic turnover rate of urine has never been analyzed mechanistically and, as further discussed, its biological meaning is mostly ascribed to the WBPT. Some considerations and assumptions need, however, to be discussed to support our biological interpretation.

First, the urinary δ^15^N kinetics post diet-switch did not only reflect the N metabolism of the animal but also that of its symbiotic rumen microbiota. Urines would theoretically also drain the previously ^15^N enriched microbiota as they renovate their own structural proteins and N pools (amino acids but also purine derivatives). However this would have quantitatively a short-term impact on urinary δ^15^N kinetics due to the high protein turnover rate of rumen microbiota (127 to 686%/d in rumen bacteria; Wallace and McPherson, 1987). Second, because of the complex and extensive urea recycling in ruminants we should not exclude the fact that the clearance rate of the previously administered ^15^N labelled urea from the whole body after stopping its administration might contribute in an undetermined extent to the urinary ^15^N depletion rate during the first hours. In this sense, it has been demonstrated in beef cattle that most infused ^15^N-labelled urea is excreted through urines within the first 48h (Wessels et al., 1997), and so its impact on urinary ^15^N depletion rate would be very short. Third, we can not exclude that differences in the N absorption, and thus in the urinary N excretion, expected across diets (Normal vs High protein diets) might contribute to the observed differences in the urinary ^15^N depletion rate. Indeed, the dietary N influx into the plasma pool, while being much smaller than the endogenous AA influx by protein degradation (Lobley, 2000), had the largest δ^15^N difference with body AA at the beginning of the diet-switch (dietary AA were largely ^15^N depleted compared to the body AA previously ^15^N enriched during the administration of ^15^N labelled urea). In consequence, the urinary δ^15^N kinetics were probably mainly driven by the dietary AA influx during the very first days (k_1_ phase), before being mostly driven by the endogenous AA influx released from the whole-body protein degradation thereafter (k_2_ phase). All these three factors (rumen microbiota metabolism, isotopic labelled urea clearance rate and differences in N absorption across treatments) would have thus a limited short impact on the urinary ^15^N turnover rate only during the first rapid phase (k1). We think that the ^15^N depletion rate in urines during the second slow phase (k2) is mostly driven by the release rate of endogenous ^15^N labeled AA from degradation of previously labeled body proteins. This progressive release of previously labeled AA will translate first into the free plasma AA pool and then into the urea-N pool resulting from their catabolism. Therefore, we assume that the urinary ^15^N depletion rate during the second slow phase (k2) is mostly the consequence of the WBPT, whereas the rate during the first rapid phase (k1) is likely a mix from the previously evoked mechanisms together with (in some undetermined extent) the protein degradation rate of some very fast animal turning-over N pools.

A limitation to our interpretation could be that some N-containing compounds in urine are metabolically unrelated to protein degradation and AA oxidation. This could represent a potential issue particularly in ruminants where urinary urea-N excretion, the main urinary nitrogenous component, may origin in a high extent from rumen ammonia production (INRA, 2018) and so from nitrogenous compounds unrelated to protein degradation and AA oxidation. It can be acknowledged that our modelled k2 likely integrates the outflow rate of those minor urinary nitrogenous compounds non related to AA oxidation and with an endogenous (animal) origin (e.g. creatine, creatinine, endogenous purine derivatives, 3-methylhistidine and other free AA). Such endogenous nitrogenous compounds were likely previously labelled during ^15^N urea administration likewise body proteins and progressively released from the animal body through the urines. However it should be stressed that i) its contribution to total urinary N remains relatively modest in high productive animals (0.05 g N/kg BW/d; INRA, 2018) and ii) for some of these nitrogenous compounds their release rate is closely related to protein turnover rate (e.g. 3-methyl-histidine as an index of muscle protein degradation and endogenous purine derivatives from nucleic acid turnover rate).

For those urinary nitrogenous compounds non related to AA oxidation and with a ruminal origin (mostly purine derivatives [allantoin and uric acid], hippuric acid and rumen ammonia mostly transformed into urea) it can be argued, as previously mentioned, that it would have probably an impact on the first rapid phase of the urinary ^15^N depletion rate (k1) but as long as the plasma absorption of such compounds remains relatively constant across time they will not have an effect on the second slow phase of this depletion (k2). Taking all this together, we can consider that even if k2 does not strictly correspond only to the WBPT, it may reflect quantitatively this flux and can be proposed as a proxy to evaluate it.

Finally, we recognize that the proposed approach based on isotope decay rates may be problematic for having a real and accurate estimation of WBPT because of the phenomenon of amino acid reutilization, (i.e. the reincorporation of labelled amino acids released by protein degradation during protein re-synthesis). Indeed, because overall rates of protein turnover markedly exceed rates of dietary protein intake much of the amino acid substrates for protein synthesis derive from protein degradation (Lobley, 2000). However, even if AA reutilization could impact the 15N values at each time point of the isotopic decay we assume that a higher protein turnover (associated also to a higher AA-reutilization) will always have a higher k2 than a low protein turnover. Thus, for comparing treatments (or even individuals) in terms of WBPT this approach can still serve as a proxy.

To evaluate the aforementioned biological interpretations, we tested the ability of our method to detect differences across two dietary factors, the protein content (Normal vs High) and AA profile (diets balanced vs unbalanced in methionine), known to impact the whole-body protein turnover rate in a different extent.

### Isotopic turnover rates across dietary factors impacting protein metabolism

Dietary protein content determines the dietary AA flux to the organism and is the major dietary determinant of the whole-body protein turnover rates in humans (Waterlow, 2006) and farm animals (Lobley, 2000) through the action of catabolic and anabolic hormones. In the present study, the urinary ^15^N-depletion rates during both the first rapid (k_1_) and second slow (k_2_) phases increased on average around 28% when increasing the metabolizable protein content by 20%. This finding is in line with the expected increases in dietary AA influx and WBPT that have been well recorded in k_1_ and k_2_, respectively. When a similar diet-switch approach was applied on adult rats fed diets at or much above protein requirements (10 vs 30 % CP), the urinary turnover rate of ^15^N increased more than two folds, in agreement with an expected higher protein turnover rate (Braun et al., 2013). Our values for the WBPT (average k2 = 9.1%/d) are, however, higher than those reported in the few studies using standard isotopic methods in growing ruminants, which themselves encompasses a wide range (3.88-8.00%/d whole-body protein FSR in growing heifers [Lobley et al., 1980]; 6.15 −7.63%/d WBPT in growing lambs [Liu et al., 1995]; calculated WBPT rate of around 1.8%/d in growing steers [Wessels et al., 1997]) but still lower than other estimates (19.7%/d WBPT rate in growing lambs [Davis et al., 1981]). Although the methods are not comparable (few days vs several months) and our higher values compared to other cattle studies (Wessels et al., 1997; Lobley et al., 1980) could be just the consequence of using late maturing breeds with high growth rates (1.70 kg/d on average for the studied period) and retaining between 35-45% of their energy as protein (Geay and Robelin, 1979), the obtained values will need further confirmation and interpretation.

We also found that the plasma protein ^15^N-depletion rate (k) increased with the dietary protein content, supporting the idea of a greater protein metabolism at all body levels as revealed through the analysis of urines. Likewise, Tsahar et al. (2008) also found in birds that the ^15^N turnover rate in plasma doubled (from around 11 to 20 %/d) when dietary protein shifted from 7 to 16 % CP. The impact of dietary protein content on the protein turnover rate of plasma proteins was demonstrated in the 50’s through the elegant study by Steinbock and Tarver (1954) who injected into rats plasma from donor rats in which the proteins were previously labelled with [^35^S] methionine. However this effect was not observed in ruminants through standard isotopic methods, where the plasma proteins FSR remained unchanged in dairy cows fed different dietary metabolizable protein content (Raggio et al., 2007) or sheep undergoing a shift between the fasted to fed state (Connell et al., 1997).

On the other hand, no changes in the rate at which urines are progressively depleted in ^15^N following an isotopic diet-switch were found between unbalanced vs balanced diets in terms of methionine content. In contrast, plasma protein ^15^N-depletion rate was slightly increased, indicating that protein FSR was enhanced for plasma proteins and likely also at other different body levels. Our findings may indicate that the observed trend for Met balanced diets to improve beef cattle performances observed in this experiment (Cantalapiedra-Hijar et al., 2017) and others (Wessels et al., 1997) could stem from an increased whole-body protein synthesis rather than a reduction in the whole-body protein degradation.

## Conclusions

We believe that the rate at which animal’s urines are depleted in ^15^N after stopping the administration of ^15^N labelled urea in the diet (i.e. isotopic turnover rate after diet switch) can be proposed as a non-invasive and simple proxy to evaluate in the long term the whole-body protein turnover rate in a high number of animals. Further studies are warranted to explore the potential and limits of this new promising isotopic approach for evaluating protein turnover. Specifically, future works should evaluate the impact of AA reutilization and urea-N recycling on the N isotopic dynamics in urine and tissues after a diet-switch.

## Acknowledgments

The authors thank Vincent Largeau and the stuff of Herbipôle for their great technical assistance during this experiment. INRA’s Phase department is acknowledged for supporting the pilot study that served to stablish the conditions of the experiment here presented.

## Supporting information

**Figure S1.**
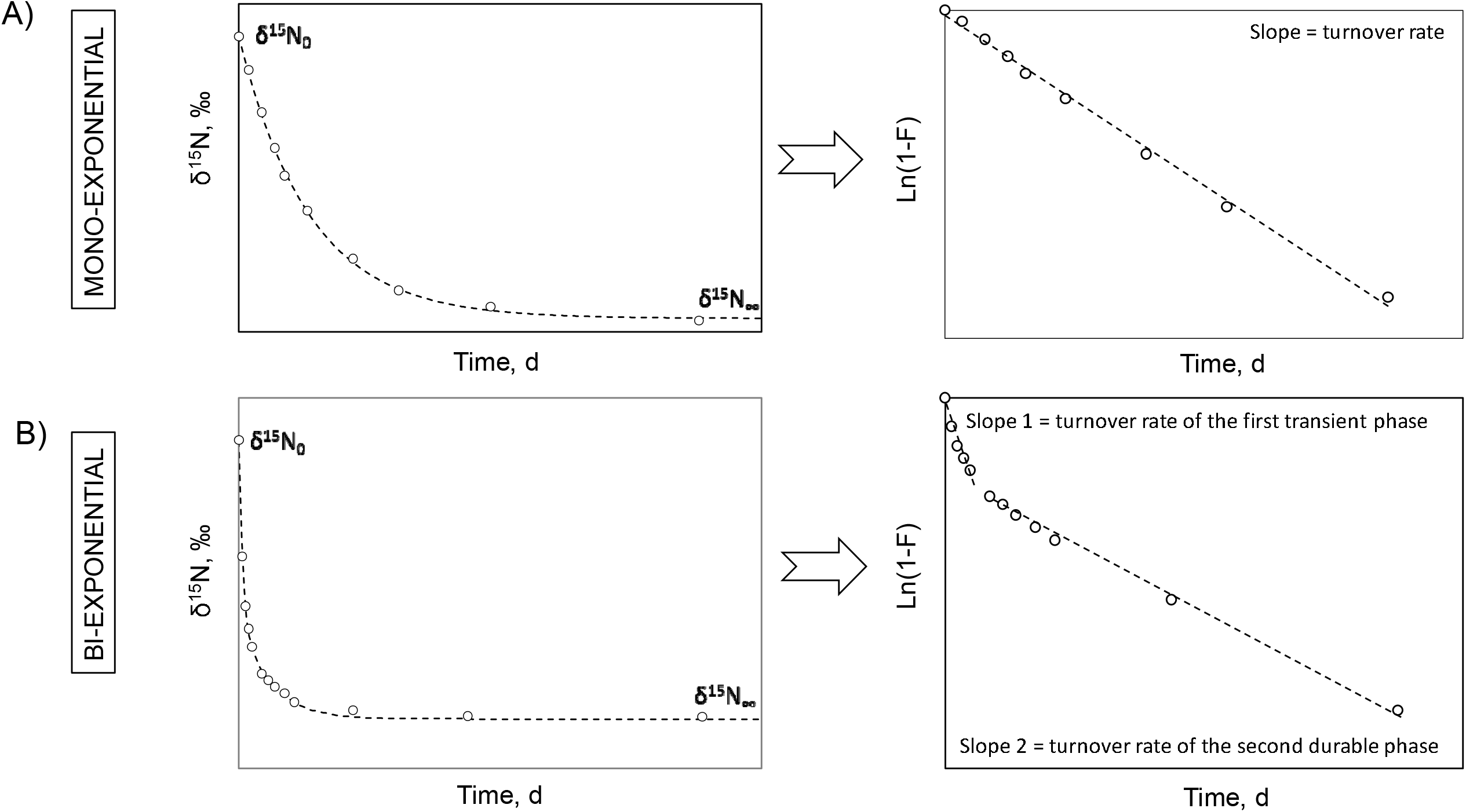
Modelling the kinetics of 15N after a 15N diet-switch in ruminants depending on whether a mono-exponential model is sufficient or a bi-exponential model is required to adequately fit the data. A) Mono-exponential model (like in plasma proteins) where δ^15^N kinetics are adequately fitted according to δ^15^N(t) = δ^15^N_∞_+ (δ^15^N_0_ - δ^15^N_∞_)×*e*^−k×t^ and a single slope is noted when the logarithm of the reaction progress variable (1-F = (δ^15^N(t) - δ^15^N_∞_)/(δ^15^N_0_ - δ^15^N_∞_)) is regressed on time. B) Bi-exponential model (like in urines) where δ^15^N kinetics are adequately fitted according to δ^15^N(t) = δ^15^N_∞_+ (δ^15^N_0_ - δ^15^N_∞_)×[p×*e*^−k_1_×t^ + (1-p)×*e*^−k_2_×t^] and two different slopes are noted when the logarithm of the reaction progress variables is regressed on time. The two different slopes represent the fractional turnover rates during a first transient (k_1_) and a second durable (k_2_) phases, respectively.

